# Genomic analysis of fast expanding bacteria reveals new molecular adaptive mechanisms

**DOI:** 10.1101/355404

**Authors:** Lars Bosshard, Stephan Peischl, Martin Ackermann, Laurent Excoffier

## Abstract

Bacterial populations have been shown to accumulate deleterious mutations during spatial expansions that overall decrease their fitness and ability to grow. However, it is unclear if and how they can respond to selection in face of this mutation load. We examine here if artificial selection can counteract the negative effects of range expansions. We investigated the molecular evolution of 20 lines (SEL) selected for fast expansions and compared them to 20 lines without artificial selection (CONTROL). We find that all 20 SEL lines have been able to increase their expansion speed relative to the ancestral line, unlike CONTROL lines, showing that enough beneficial mutations are produced during spatial expansions to counteract the negative effect of expansion load. Importantly, SEL and CONTROL lines have similar numbers of mutations indicating that they evolved for the same number of generations and that increased fitness is not due to a purging of deleterious mutations. We find that loss of function (LOF) mutations are better at explaining the increased expansion speed of SEL lines than non-synonymous mutations or a combination of the two. Interestingly, most LOF mutations are found in simple sequence repeats located in genes involved in gene regulation and gene expression. We postulate that such potentially reversible mutations could play a major role in the rapid adaptation of bacteria to changing environmental conditions by shutting down expensive genes and adjusting gene expression.

**Author Summary:** We investigated if strong artificial selection for fast expansion can counteract the negative effects of range expansion which had been shown to lead to an accumulation of deleterious mutations. This experiments showed that i) an increase in expansion speed could occur if bacteria were selected from the largest protruding sectors, and ii) that artificially selected bacterial lines accumulated about the same number of mutations than simply expanding line suggesting that the observed increased fitness is not due to increased purifying selection where deleterious mutations would have been removed in fast growing lines. We find that loss of function (LOF) mutations are best explaining the observed increased expansion speed in selected lines. These mutations, which are known to play an important role in adaptive processes in bacterial populations, frequently consist in small insertion-deletions in simple sequence repeats, and are thus relatively easily reversible. They could thus act as switches that can reversibly shut down genes. Our results therefore suggest that shutting down expensive genes and adjusting gene expression are important for adaptive processes during range expansion.

## Introduction

Theoretical studies have recently predicted that spatial expansion of populations can lead to the fixation of deleterious mutations (1, 2) due to small effective size and inefficient selection on range margins. When a spatial expansion proceeds for a long time, edge populations tend to accumulate a series of deleterious mutations, leading to a decrease in fitness over time and space (3, 4). This “expansion load” (3) can potentially affect the speed of the expansion and impose constraints on the limits of a species range (5). Our recent empirical work supported these theoretical predictions, and showed that spatially expanding bacterial colonies accumulated deleterious mutations that impacted their fitness (6).

However, the accumulation of deleterious mutations is potentially not the only relevant evolutionary process in expanding populations. Under some conditions, populations that expand their range might experience selection for rapid range expansion (7). There are potentially two different mechanisms for adaption for individuals at the front in a radial expansion: (i) Resources are more abundant on the front than in the core of the colony therefore cells of the same type grow faster at the front than in the core and adaption can occur by enable individuals to reach the front. (ii) Fast expanding bacteria at the front can outcompete bacteria expanding more slowly in neighboring sectors and thus expand sideways. While reaching the front does not necessarily lead to a faster expansion since there can be a negative tradeoff between growth rate and the ability to reach the front, competition at the front should increase the expansion speed in all cases. In general, selection for rapid range expansion can occur under the same conditions as selection for rapid dispersal, with the caveat that selection for rapid range expansion requires more stringent conditions. Indeed, it requires that different groups of individuals compete with each other (groups that are located at different locations of a two dimensional expansion front), in contrast to evolution for rapid dispersal that is based on competition between different individuals competing for being at the front. One would thus expect that selection for rapid range expansion is only effective in certain types of organisms, namely organisms that form very large populations and expand their ranges on wide fronts where different clonal sectors can compete.

Under conditions where selection for rapid range expansion is expected, it is important to consider its consequences for genome evolution. As mentioned above, range expansion leads to a reduction in the effective population size on the front and consequently to an accumulation of deleterious mutations in edge individuals. It is yet unclear how this mutation accumulation process would interfere and potentially hinder adaptive genome evolution, if for instance there is (artificial) selection for rapid range expansion, One can imagine two fundamentally different, but not mutually exclusive, ways how mutation accumulation and adaptive evolution can interact. First, the fastest expanding groups of individual could harbor fewer deleterious mutations than other groups at the expansion front and thus grow faster. Second, the fastest expanding groups of individuals could accumulate similar amounts of deleterious mutations as other groups of individuals, but they could harbor more beneficial mutations, or an equal number of positively selected mutations that would have larger effects. Discriminating between these different scenarios is a novel and interesting endeavor.

Here, we addressed this question by performing an evolution experiment with populations of the bacterium *E. coli*. We let replicated populations of this bacterium expand their range by placing them on a solid surface of nutritious medium and letting them expand radially, forming an approximately circular expanding population. After three days of expansion (corresponding to about 127 generations), we selected the section of the colony edge that had expanded furthest. We collected about one million individuals from the outer edge of this protruding sector and transferred them to a new habitat where we let them again expand. We thus imposed a regime where only individuals belonging to the fastest growing sector could continue to evolve, whereas all other individuals were removed.

We performed this evolution experiment independently in 20 populations, starting from the same ancestral strain. In addition, we also evolved 20 control populations that were propagated in the same way with the important difference that the sector of the front from which individuals were selected to be transferred to a new habitat was chosen at random, thus without imposing any selection for rapid range expansion. As in our previous range expansion experiment, we worked with a mutator strain of *E. coli* having a mutation rate about 200 times higher than that of wild-type *E. coli*.

Our goal here is two-fold. First, we ask whether there is a response to selection for increased range expansion. As mentioned above, it is a priori not clear how the balance between mutation accumulation and adaptive evolution will occur in such an experiment. As a consequence, it is not clear whether this regime allows the selection of an increased expansion rate. Our second goal is to analyze the magnitude and the quality of the genomic changes under control and selected conditions. We are thus interested in examining how the interplay between mutation accumulation and adaptive evolution shapes the genomes of the populations that we selected for rapid expansion, and how this compares to the genomic evolution of controls without selection.

If we were to observe more rapid range expansions, we could ask more specifically which biological alterations would underlie such a response. One possibility to increase expansion speed is to increase the rate at which individual cells grow and divide. This would likely involve mutations in metabolic pathways, genes for nutrient transporter, and genes responsible for gene expression regulation. Another possible mechanism could be spatial sorting (8), where bacteria would evolve phenotypic traits that would allow them to move within the expanding populations and reach the edge of the expansion faster, without necessarily having a higher growth rate. This spatial sorting phenomenon has been invoked in a recent study where it was shown that alterations of surface proteins influenced the positioning of bacterial cells within an expanding population (9). The dissection of newly accumulated mutations should thus provide us with useful insights into the molecular bases of adaptation during range expansions.

## Results

### Increase in expansion speed

We let *E. coli* strains expand radially on top of agar plates for 13 periods of 3 days. We compared 20 lines that were sampled at a random place after each period of 3 days (CONTROL lines) to 20 lines that were sampled at the point of the colony that expanded the farthest (SEL lines) (see Methods). The colony size was measured after every growth period of 3 days. We find that the CONTROL colony sizes decreased significantly over time (−77 μm/day, 95% C.I. [-95;-60], p-value: < 2 × 10^−16^), whereas the size of the SEL colonies increased significantly (227μm/day, 95% C.I. [192; 262], p-value: < 2 × 10^−16^) (**Figure 1**). Thus, after 39 days, CONTROL colony sizes decreased by 33% on average (t-test: p-value <2.2 10 ^−16^, 95% C.I. [29; 38]), whereas those of SEL lines increased by 130% (t-test: p-value <2.2 10 ^−16^, 95% C.I. [119; 142]). We used a linear mixed-effect model to determine the dynamics of expansion velocity change over time. A quadratic term in the mixed effect model explains the data significantly better than a simple linear model (CONTROL: likelihood ratio 7.14, p-value = 0.0075; SEL: likelihood ratio 25.98, p-value < 0.0001). There is thus a saturation effect in both conditions (SEL and CONTROL) over time.

**Figure 1:**
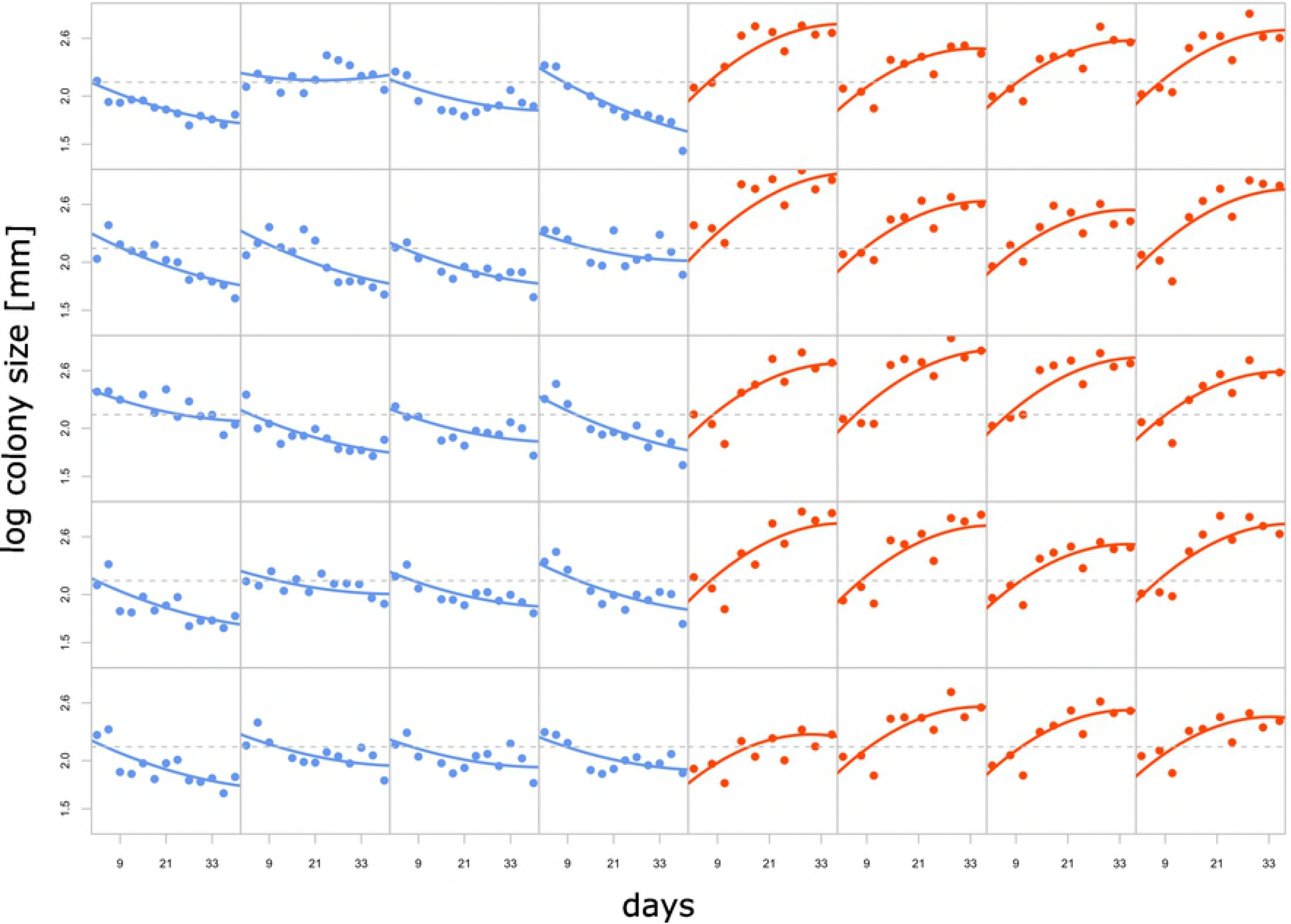
Evolution of colony size after 3 days of growth on LB agar plates. Blue: CONTROL lines, red: SEL lines. The x axis represents the total days of evolution. The horizontal dashed lines represent the average colony size measured in the first 3 days of the experiment over all SEL and CONTROL lines. Solid lines represent the line specific regression lines from the mixed-effect regression analysis.

### Similar number of mutations in both lines

The average number of mutations is 124.9 in CONTROL lines and 129.8 in SEL lines (**Figure 2**). We tested for a significant difference in mutations numbers between the groups using a non-parametric Mann-Whitney test, since the variance in the number of mutations was found significantly different in the two groups (variance SEL = 1273.36, variance CONTROL = 499.15, p-value: 0.048 with a Bartlett test of homogeneity of variance). However, we do not find any significant difference in the number of accumulated mutations between the SEL and CONTROL lines (p-value: 0.7048). Similarly, the dN/dS ratios were not significantly different between the two groups (CONTROL lines: 1.138, SEL lines: 0.995, Mann-Whitney test: p-value =0.097). As previously described (6), the mutations are distributed along the genome with a periodic pattern that is repeated nearly in mirror-image across the genome (**Figure 2B**) centered on the origin of the genome replication. This uneven genomic distribution of the mutations implies that there is a variable mutation rate during the replication of the genome, but that the two replication forks have similar changes in mutation rate as they traverse the chromosome. We estimated these variable mutation rates across the genome by a wavelet transformation (10) (**Figure 2B**).

**Figure 2:**
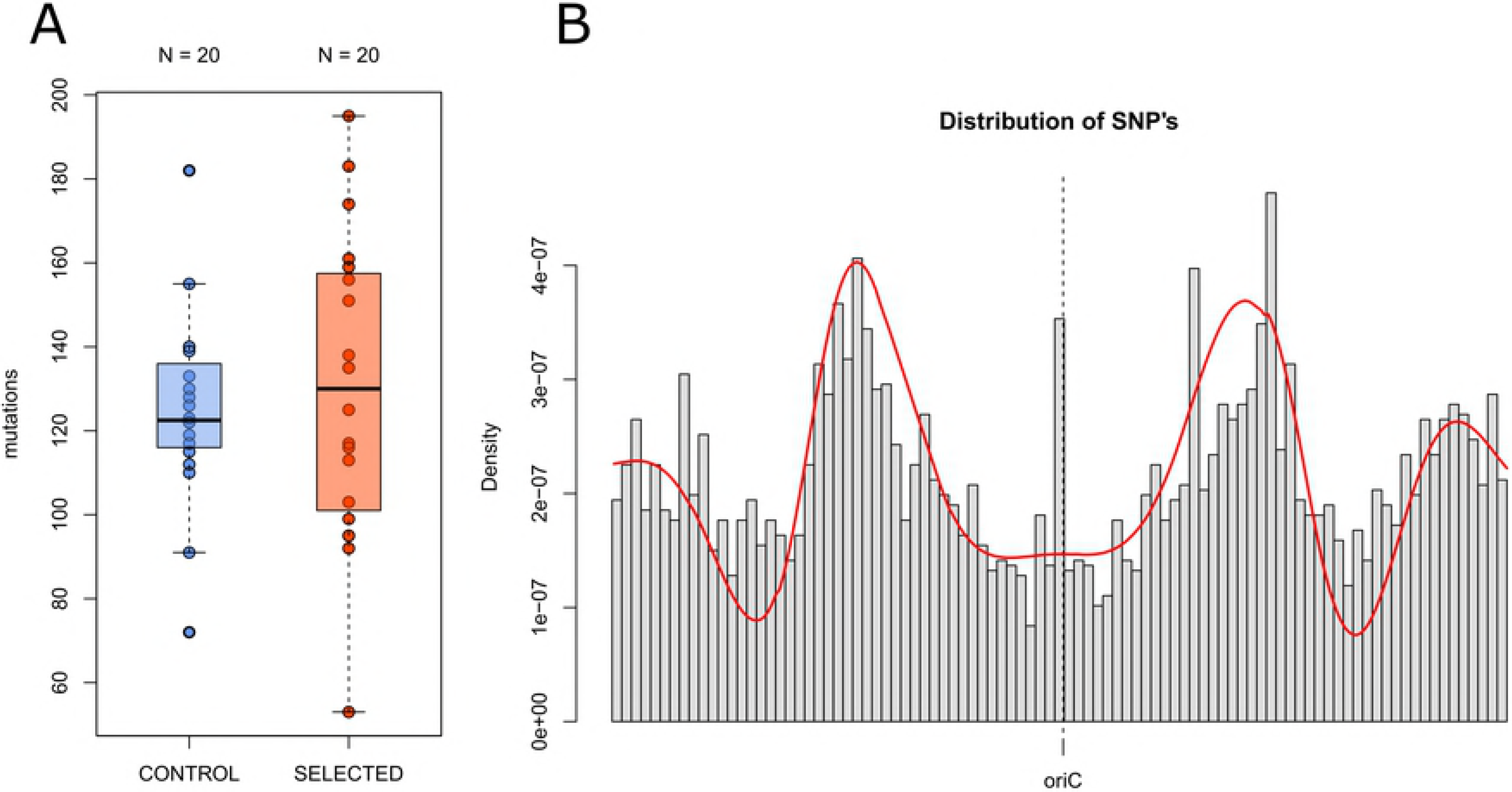
**A)** Number of mutations after 39 days of expansion blue: CONTROL lines, red: SELECTED lines. The variance in the number of mutations is significantly different (Bartlett test of homogeneity of variance: p-value= 0.048) **B)** Distribution of SEL and CONTROL mutations along the genome. The origin of the genome replication is indicated as oriC, red line: Wavelet transformation of the distribution of the mutations

### LOF mutations as a main driver of adaptation

We used Elastic Net (EN) regression (11), which performs both variable selection and variable regularization (see Material and Methods), to determine the subset of genes that have the largest effect on the expansion speed in bacteria. The resulting significant coefficient associated to a gene is its net effect on final colony size relative to the initial colony size. With this analysis, we determined which genes explain the difference in colony size between the SEL and CONTROL conditions. We used the EN regression to predict colony size from three different sets of mutations. First, we used the combination of all non-synonymous substitutions, as well as frameshift and non-sense mutations (**Table S1**). Second, we analyzed separately frameshift and non-sense mutations. Note that non-sense mutations can be considered loss of function (LOF) mutations for a specific gene (**Table 1**). Finally, we used only non-synonymous mutations, which could be a target for adaptation without loss of gene function (**Table S2**). We then compared the mean cross-validation error of these models, and find that LOF mutations significantly better explain colony size change (mean error=0.3258) than all mutations taken together (mean error = 0.4105, p = 1.06 10^−10^) or than non-synonymous mutation alone (mean error = 0.4033, p = 4.85 10^−10^).

**Table 1:**
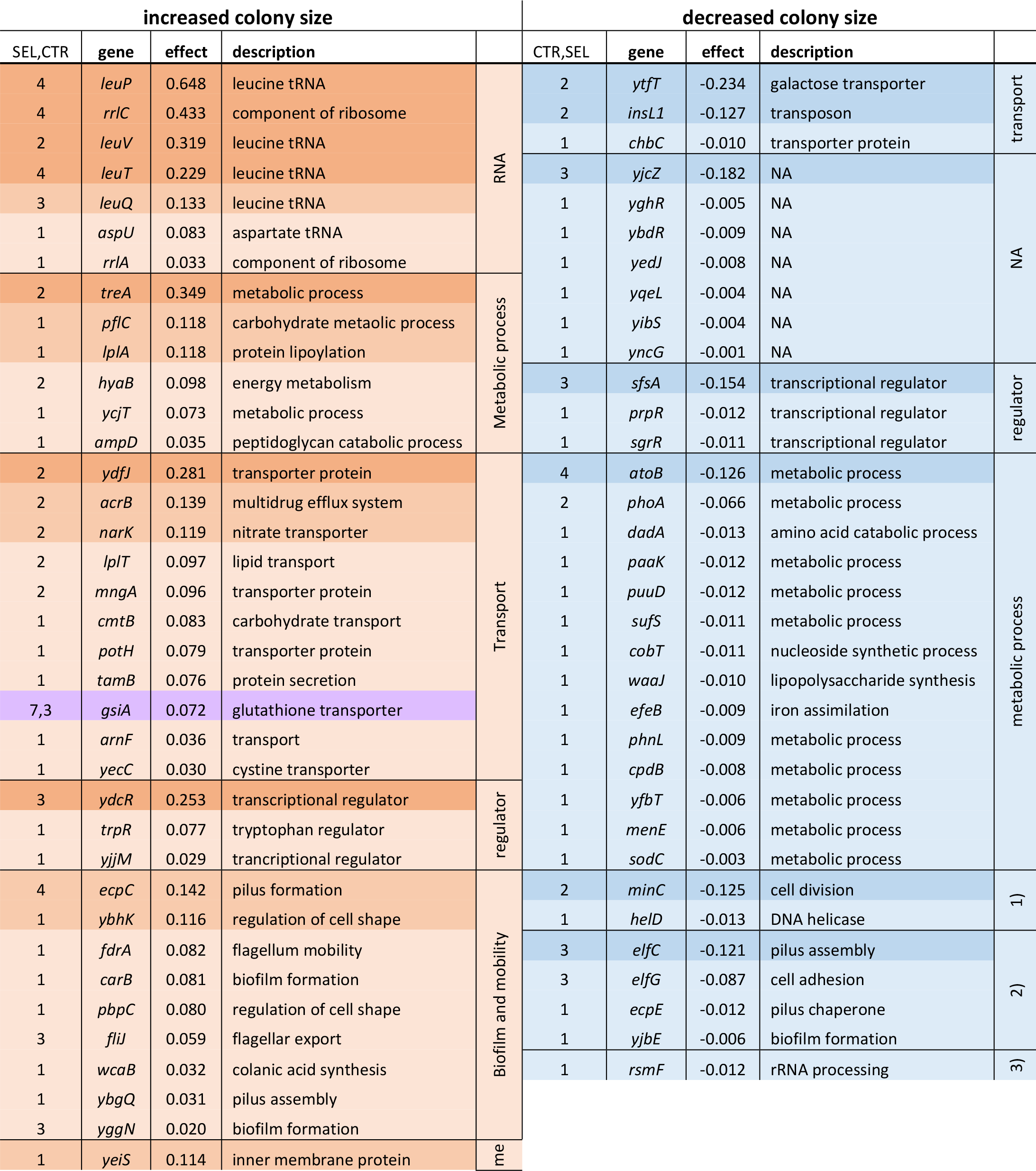
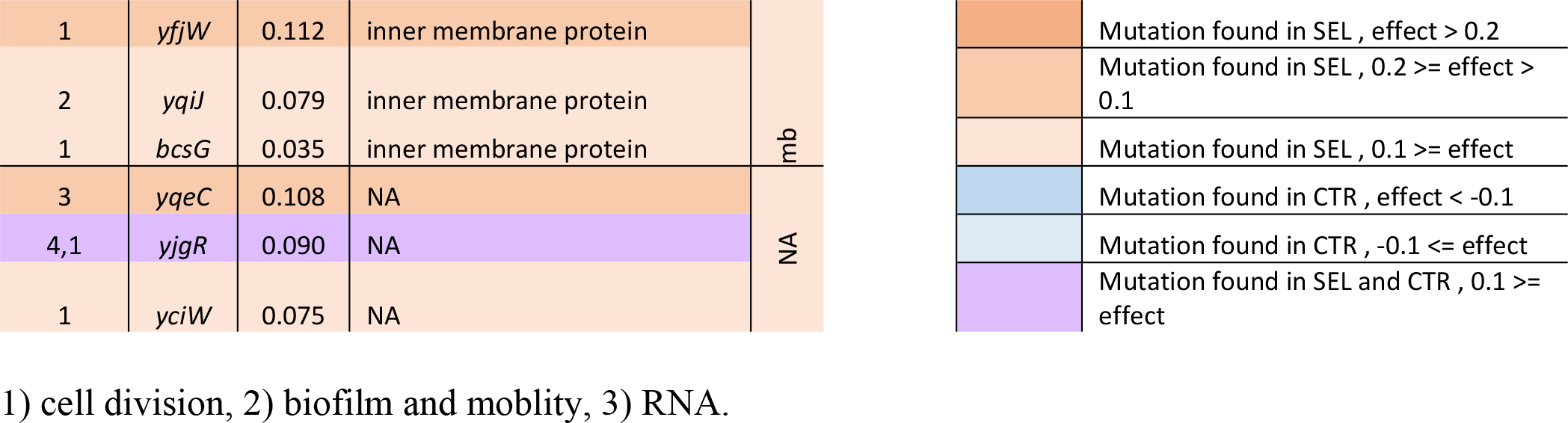
Effects of LOF mutations on bacterial growth, as inferred by the Elastic Net regression. Effect sizes are relative to the initial colony size. Orange: genes that have mutations in SEL lines, blue: genes that have mutations in CONTROL lines, violet: genes that have mutations in both CONTROL and SEL lines.

Focusing on LOF mutations, we find a total of 43 genes significantly associated with increased colony size and 34 genes significantly associated with a colony size reduction. Quite remarkably, almost all genes leading to a significant increase in colony size are targets of mutations in SEL lines, whereas all genes leading to a significant decrease in colony size are target of mutations in CONTROL lines (**Table 1**). The only exceptions are two genes connected to ATPases (*gsiA* and *yjgR*), where mutations occur in both SEL and CONTROL line. Mutations in these genes lead to an increased colony size, in agreement with the previous observation that there is still some adaptation going on in CONTROL lines (6).

Genes leading to either increased or decreased colony size are involved in metabolic process, transport, gene regulation, biofilm formation, as well as tRNA and rRNA genes. There is evidence for an association between the number of significant genes we find in the gene categories and the impact that mutation in these significant genes have on colony size (increase or decrease of colony size) (Chi-squared test, p-value = 0.009) (**Figure S5**). Compared to genes that decrease colony size, there are more genes leading to an increase in colony size that are involved in the transport of substances through the cell membrane or in processes associated with ribosomes and in tRNAs. In contrast, there are more genes that lead to a decrease in colony size that are involved in metabolic processes (**Table 1**, **Figure S5**). Additionally, we find two genes where mutations lead to a decreased in colony size that are involved in cell division. Note however, that a separate GO enrichment analysis for genes leading to significant increase or decreased colony size with the EN analysis did not reveal any significant term.

Finally, it is worth emphasizing that seven genes leading to an increased colony expansion are connected to tRNAs (*leuP, leuV, leuT, leuQ, aspU*) or rRNAs (*rrlA, rrlC*), but that only one gene connected to rRNA leads to a decrease in colony expansion (*rsmF*) (**Table 1**). Quite remarkably, four out of eight copies of the leucine tRNA have been targeted by indel mutations and all lead to a large increase in colony size.

### Convergent Adaptation via LOF Mutations

We looked for signals of convergent adaptation by searching for mutations that have targeted the same gene in unrelated lines, which is usually taken as evidence for a signal of adaptive processes (12). We therefore tested if some genes were targeted by the 3044 observed non-synonymous and LOF mutations more frequently than expected by chance. We simulated the random occurrence of 3044 mutations along the genome, taking explicitly into account the differential mutation rates across the genome as inferred in **Figure 2B**, and we compared the simulated and observed numbers of genes targeted by these mutations (**Figure 3**). Non-synonymous and LOF mutations were analysed separately, and we categorized the genes in three groups: genes with at least one mutation in either i) CONTROL lines, ii) SEL lines, and iii) genes that mutated in both SEL and CONTROL lines. The analysis of non-synonymous mutations shows no departure from expectations in any category (**Figure 3A**), whereas we find more genes jointly targeted by LOF mutations between CONTROL and SEL line than expected (**Figure 3B**). Note that this excess might be due to hotspots for frameshift mutations like single sequence repeats (SSR) regions in the genome (13). More interestingly, we observe significantly fewer genes than expected to have been targeted by LOF mutations in SEL lines. In other words, LOF mutations are more clustered than expected by chance in SEL lines. We indeed find that there is a significant excess of genes that have been the target of 2 and of 3 or more LOF mutations in SEL lines (**Figure 3D**) and of 3 or more LOF mutations in CONTROL (**Figure 3F**). Note however, that there is no deviation from expected counts of mutations per gene for non-synonymous mutation (**Figure 3C, 3E**), such that only LOF mutations seem to preferentially accumulate in specific genes.

**Figure 3:**
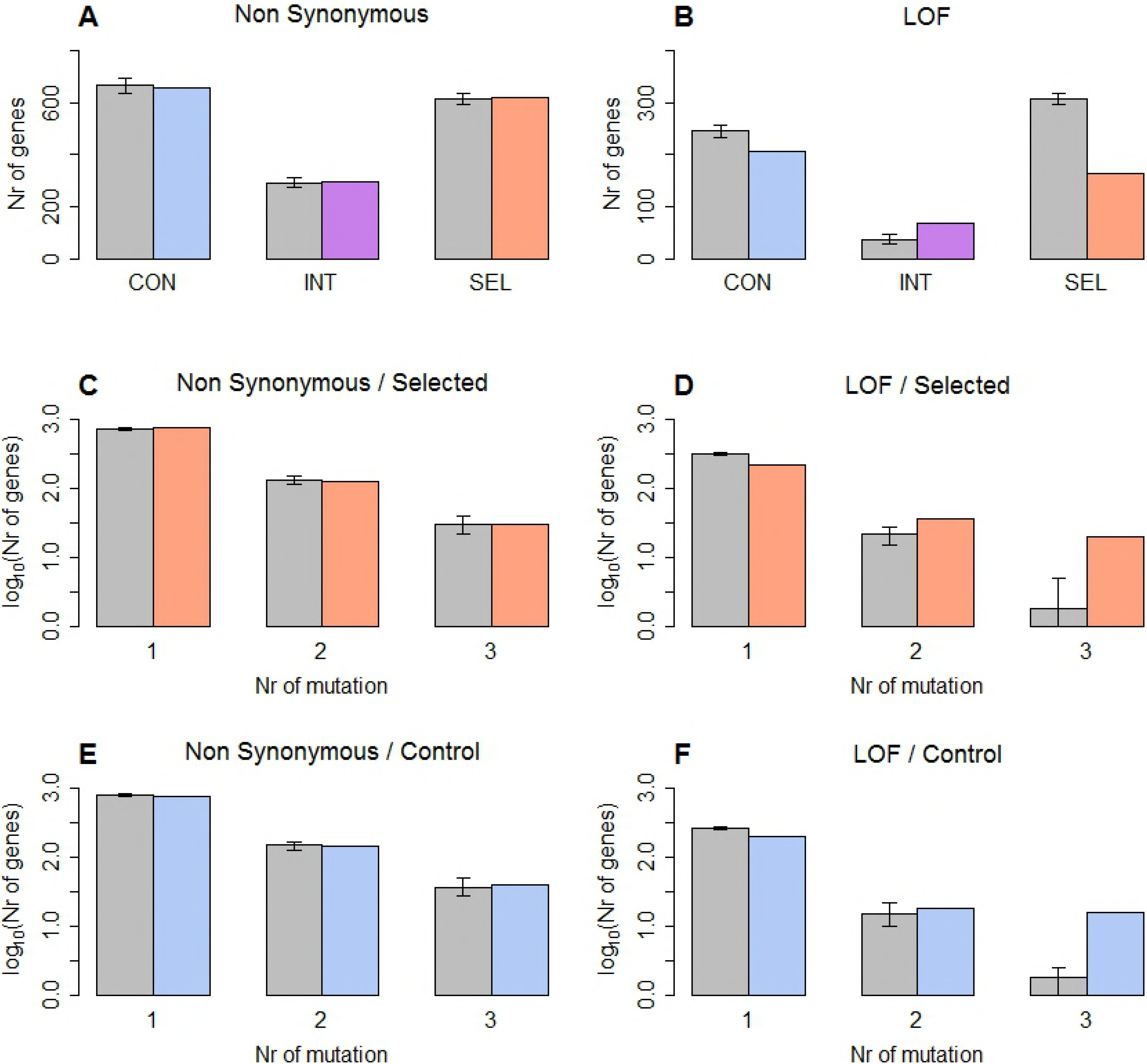
Distribution of mutations among genes. **A, B:** Number of genes with at least one mutation in either CONTROL or SEL lines, and number of genes that mutated in the SEL and the CONTROL experiment (INT). Whiskers indicate limits of empirical 95% CI computed from 1000 simulations. Grey bars: expected numbers. Color bars: observed numbers **A:** non-synonymous mutations **B:** LOF mutation. **C-F:** Number of genes with a given number of mutations in SEL and CONTROL experiments. **C, E:** Genes with non synonymous mutations, **D, F:** Genes with LOF mutation. Blue: CONTROL lines, orange: SEL lines, violet: CONTROL lines and SEL lines

### Enrichment of non-synonymous and LOF mutations in flagella genes

Since there is evidence that mutations in SEL and CONTROL lines are more clustered than expected, we then looked if there were any GO terms with significantly enriched numbers of genes that have non-synonymous substitutions, frameshift mutations, or non-sense mutations. We found 3 significant GO terms in SEL lines: taxis GO:0042330 (q-value = 1.58 10^−4^), amine catabolic process GO:0009310 (q-value = 0.04), colonic acid biosynthetic process GO:0009242 (q-value = 0.04), and one significant GO terms in CONTROL lines: taxis GO:0042330 (q-value = 0.004) (Table S3). Interestingly, the genes showing non-synonymous substitutions or LOF mutation enrichment in our top GO term “taxis” are all involved in the formation of the flagella (*fliG*, *fliM, fliN, fliO, motA, motB, fliJ*), and support the view that function of flagella is hindered by these mutations. Indeed, several genes hit by non-synonymous substitutions or LOF mutations are actively involved in the growth and assembly of the flagella. Nine genes have mutations in both SEL and CONTROL groups, like the flagella hook protein *FlgE*, the protein controlling flagella hook-substructure *FliK,* a protein that is involved in the assembly of the flagellar motor *FlgG*, the protein which makes up the peptidoglycan ring of the flagellar basal body *FlgI*, one of the components of the flagellar motor’s switch complex *FliM*, three components of the flagellar export apparatus *FliI, FliO,* and *FliP*, and an element of the flagellar motor complex *MotA* (**Table S4**). Focusing more generally on the 36 genes involved in the formation of the flagella, we found 13 non-synonymous mutations, 3 synonymous mutations, 9 frameshifts in the CONTROL lines and 11 non-synonymous mutations, 2 synonymous mutations, and 13 frameshifts in the SEL lines (**Table S4**), suggesting the occurrence of non- or sub-functional flagella in 15 out of 20 CONTROL lines and in 16 out of 20 SEL lines. In line with this observation, a motility test reveals that 14/20 CONTROL lines and all 20 SEL lines have a reduced motility as compared to the ancestor line (**Figure S5**). Moreover, 8/20 CONTROL lines and 11/20 SEL lines have LOF mutations in flagella genes, but there is no significant correlation between motility and number of LOF mutations in these genes (p-value=0.3358).

Although there is no strong signal of adaptation when considering the dN/dS ratio at the whole genome level (dN/dS = 1.06), we found quite high dN/dS ratios in flagella genes (CONTROL: dN/dS = 2.369 [13 non-synonymous, 3 synonymous] p-value = 0.20; SEL: dN/dS = 3.007 [11 non-synonymous, 2 synonymous] p-value = 0.16). Even though these ratios are not significant when analyzed separately in SEL and CONTROL lines due to the small total number of mutations, the dN/dS ratio is significant if we pool the two lines (dN/dS = 2.62, p-value = 0.05). These results suggest the occurrence of adaptive non-synonymous mutations in both conditions (SEL and CONTROL), and are in line with the previous observation that the colony size of CONTROL lines has initially increased before steadily decreasing over time (6). It appears that selection for flagella loss also occurred in SEL lines, but in addition, our results suggest that the LB medium is better exploited for the use of amino acids (amine catabolic process; GO:0009310) and that there is also a potential adaption to the environmental conditions in SEL lines by modifying the synthesis of the colanic acid (colonic acid biosynthetic process; GO:0009242).

## Discussion

We compared *E. coli* colonies that expanded naturally on agar plates to colonies that were selected for fast expansion to see if the latter selection regime could counterbalance the accumulation of deleterious mutations occurring on the edge of the colonies during range expansions (6), and if yes, determine which molecular mechanisms were involved in that process. Indeed, we find that all 20 SEL lines that were repeatedly selected for being the furthest from their inoculation point have been able to increase their expansion speed over time, unlike most CONTROL lines (18/20), which showed a decreased expansion speed. Thus, despite having grown under the same conditions as the CONTROL lines, a simple form of artificial selection has allowed SEL lines to increase colony size by130% compared to the ancestral line, corresponding to about 10% speed gain between each transfer.

Interestingly, SEL lines have on average the same number of mutations as CONTROL lines (**Figure 3**), implying that i) they have evolved for the same number of generations and ii) their fitness gain is not due to the purging of deleterious mutations or a selection for fewer harmful mutations, but that it is rather due to the selection of particularly beneficial mutations. Indeed, selection in SEL lines is imposed only during the transfer of the sample to a new plate (every ~ 127 generations), such that drift on the edge of the colonies should be as strong as in CONTROL lines. Therefore, the speed gain observed in SEL lines is due to the selection of clones present in fast growing sectors that overtook slow growing sectors where deleterious mutations were present. Indeed a small disadvantage of some sectors can lead to their loss on the expanding colony front, which tends to be occupied by the fastest growing colonies (14). The selection regime we have imposed thus seems to be akin to artificial spatial sorting (8), by which phenotypes with faster dispersal abilities colonizing the front are positively selected. Note that we observe however a saturation effect over time in both SEL and CONTROL lines, which is in line with Fisher’s Geometric model (15, 16) predicting that the proportion of beneficial mutations entering a population, and hence the potential for adaption, is decreasing with the population’s distance to the optimum (17). When the SEL lines get closer to their optimum, the proportion of new beneficial mutations is thus expected to decrease and hence the speed of adaptation goes down. Conversely, by accumulating deleterious mutations, the CONTROL lines are expected to have their fitness decrease and thus to be further from their optimum, allowing for a higher influx of beneficial mutations, and at the same time a decreasing expansion speed will allow selection to become more efficient and to better purge deleterious mutations (4).

We used the Elastic Net framework to evidence selection by finding genes where mutations had a significant impact on growth and by quantifying the effect size of these mutations in different genes. Interestingly, LOF mutations (frameshift and non-sense mutations) better explain colony size change than non-synonymous mutations or than considering LOF and non-synonymous mutations altogether. The average positive effect of mutations from the Elastic Net analysis is significantly larger than the average negative effect (p-value = 1.03×10^−7^) (**Figure S4**). It implies that the beneficial mutation identified by the Elastic Net analysis have on average a greater impact on the colony size than the deleterious mutations, which makes sense since the mean fitness gain of SEL lines is larger than the fitness loss of CONTROL lines, despite both lines having accumulated about the same number of mutations.

Additionally, we find that LOF mutations have preferentially targeted a restricted set of genes in both CONTROL and SEL lines (**Figure 3B**), which is potentially due to hotspots for frameshift mutations in the genome like single sequence repeats (SSR). In agreement with this hypothesis, we find that 84% (CONTROL) and 86% (SEL) of frameshift mutations are in SSR regions that are known to be hypermutable sites (13). However, LOF mutations in the SEL lines are clustered in fewer genes than in the CONTROL lines suggesting they could still be the signature of convergent adaptive events. Several studies have proposed that SSR mutations could act as a “switch” that can shut down genes (13, 18, 19). Interestingly, these frameshift LOF mutations in mononucleotide repeat regions are reversible, since back mutations can restore the original reading frame. Thus, this mechanism could provide an important source of variation that would increase the adaptative potential of bacterial populations. Worth of note, we find a significant enrichment of genes with high SSR density in GO terms that are involved in gene regulation and gene expression (**Table S5**). On the other hand, genes that have no SSR are enriched in GO terms involved in programmed cell death or other essential cellular and DNA replication functions such as transcription, translation, or protein disassembly (**Table S6**). These GO results suggest that there has been selection against SSRs in essential genes and positive selection for SSRs in regulatory genes. The combination of a high mutation rate and the possibility of back mutations in SSRs close to regulatory genes could lead to a faster adaptation to new conditions by modifying gene expression of non-essential genes, which would imply that the location of SSRs might have been under selection in *E. coli*.

Quite unexpectedly, we find that 4 out of 8 genes copies of the leucine tRNA have frameshift mutations that are associated to increased colony size (leuP, leuV, leuQ, and leuT) (**Table 1**). These tRNAs are all targeting the CUG codon, which is one of the most abundant codon used by *E. coli*. There are in total four copies of these tRNAs and there is not more than one mutation in these genes per line, implying that there is still a functional version of this tRNA present in each cell. It is likely that the mutations alter the structure of the tRNA, which in turn could affect the affinity of the tRNA to the amino acid. Interestingly, all 13 mutations found in leucine tRNA genes are in the acceptor stem that connects the tRNA to the amino acid. It has been shown that besides the anticodon loop the acceptor stem part is also important for the recognition of the amino acid (20). Altering the affinity of one of the 8 leucine tRNA copies could change bacterial fitness by changing the proportion of tRNAs that are connected to leucine since there would be a lower amount of functional tRNAs for leucine (21). Mutations in these tRNAs could thus optimize the level of rare and common leucine tRNAs that are charged in the cells since it could lead to non-functional tRNAs and hence to a lower leucine tRNA level compared to other tRNAs. This suggests that these modified tRNAs could speed up protein production in SEL lines.

Also of interest, we find that non-synonymous mutations in the RNA polymerase (rpoC) lead to an important increase in colony size of SEL lines (**Table S3**), in line with previous studies showing that mutations in the RNA polymerase gene were adaptive for optimal growth (12, 22). rpoC is part of the β’ subunit of RNA polymerase, which is involved in the enzymatic function of the polymerase, especially at the promoter melting stage, and 4 out of 20 SEL strains show non-synonymous mutations in the β’ rpoC subunit. In SEL lines, we found an enrichment for mutations in the GO term amine catabolic process (GO:0009310), which suggests that there has been an adaptation to the LB medium since this medium is rich in oligopeptides. *E. coli* indeed has several oligopeptidases and peptidases, enabling it to recover free amino acids from many oligopeptides (23). There is also an enrichment of non-synonymous and LOF mutations in colanic acid biosynthetic genes in SEL lines (GO:0009242). Colanic acid is a negatively charged polymer of glucose, galactose, fructose, and glucuronic acid that forms a protective capsule surrounding the bacterial cell surface. Previous studies have shown that colanic acid synthesis is upregulated in biofilms and that it has a potential protective function in hostile environments (24, 25). Alternatively, since the production of these genes is costly, loss of function mutations could lower these costs (26), potentially allowing bacteria to grow faster.

Both CONTROL lines and SEL lines have more mutations suggestive of adaptation in the taxis GO term (GO:0042330). This potentially adaptive mutations could explain the initial fitness increase of the CONTROL lines that was described previously (6). The mutations connected to the taxis GO term are in genes that are components of the flagellum. LOF mutations in flagella genes could lower the cost of production of flagella. In addition, bacteria with deficient or absent flagella could more easily invade the edge of the colony due to a cell sorting mechanism like that recently shown for bacteria with lower pili density (9). In that case, flagella-deficient bacteria would not have any particular growth or reproductive advantage, but they would just more easily disperse and invade the wave front. However, recent studies have shown that flagella has an architectural function within biofilms, as they are expressed at the front of the bacterial colony and tether cells together in a mesh (27). All SEL lines and 14/20 CONTROL lines show a reduced flagellum functionality, but not a complete loss of motility. Further studies would thus be necessary to investigate the exact impact of altered flagella genes on the colony structure and expansion speed.

LOF mutations are generally considered as important for adaptive processes in bacteria (26, 28), and we provide here evidence that they play a key role in the increased expansion speed of SEL lines. We observed many SSR LOF mutation in genes involved in gene regulation and gene expression, as well as several non-synonymous mutations in the RNA polymerase gene that is also influencing gene regulation. It suggests that the observed differences between SEL and CONTROL lines are mainly due to a combination of shutting down expensive genes and modified gene expression leading to faster growth. Overall, our detailed genomic analyses, which have allowed us to reveal the exact genetic mechanisms involved in the control of bacterial expansions, suggest that SSR LOF mutations could have been selected to act as reversible gene switches and expression modulator, allowing bacteria to quickly adapt to a variety of situations that extend much beyond mere range expansions.

## Methods

### Bacterial strain

We used *E. coli* K12 MG 1655 mutator strains where the expression of the *mutS* gene is directly controlled by the arabinose promoter *pBAD* inserted in front of the *mutS* gene. In absence of arabinose, *mutS* is not expressed, leading to a higher spontaneous mutation rate due to the inactivation of the methyl-directed mismatch repair system (29). Bacteria grown in presence of arabinose express the *mutS* gene and thus have a lower spontaneous mutation rate. This feature was used to prevent accumulation of mutations during bacterial manipulations performed outside evolutionary experiments. Additionally, our strain includes a GFP marker located in the lac operon, which can be induced by Isopropyl β-D-1-thiogalactopyranoside (IPTG).

### Experimental setup

Bacterial strains were grown on LB agar plates at 37°C for a total duration of 39 days. The strains were transferred on new agar plates every 3 days. An image of the colony was taken before transferring the strains to a new plate. At each transfer, 100 million cells were sampled from the colony front using a sterile pipette tip and resuspended in 100 μl 0.85% NaCl solution. One million cells were then used to inoculate a new plate (**Figure 1A**). We used two different sampling protocols. 1) No selection (CONTROL): we drew a line from the centre of the plate to the edge before the plate was inoculated, and after three days of growth, bacteria were sampled at the point of contact of the line and the colony. 2) Artificial selection (SEL): we let the bacteria grow for 3 days, and the edge of the furthest expanding sector of the colony was sampled (**Figure 4B**). The artificial selection experiment was performed on 20 SEL lines, which were then compared to 20 CONTROL lines randomly selected from a previous experiment (6). In both cases, 13 transfers were performed for each line in a period of 39 days (**Figure 4C**).

**Figure 4:**
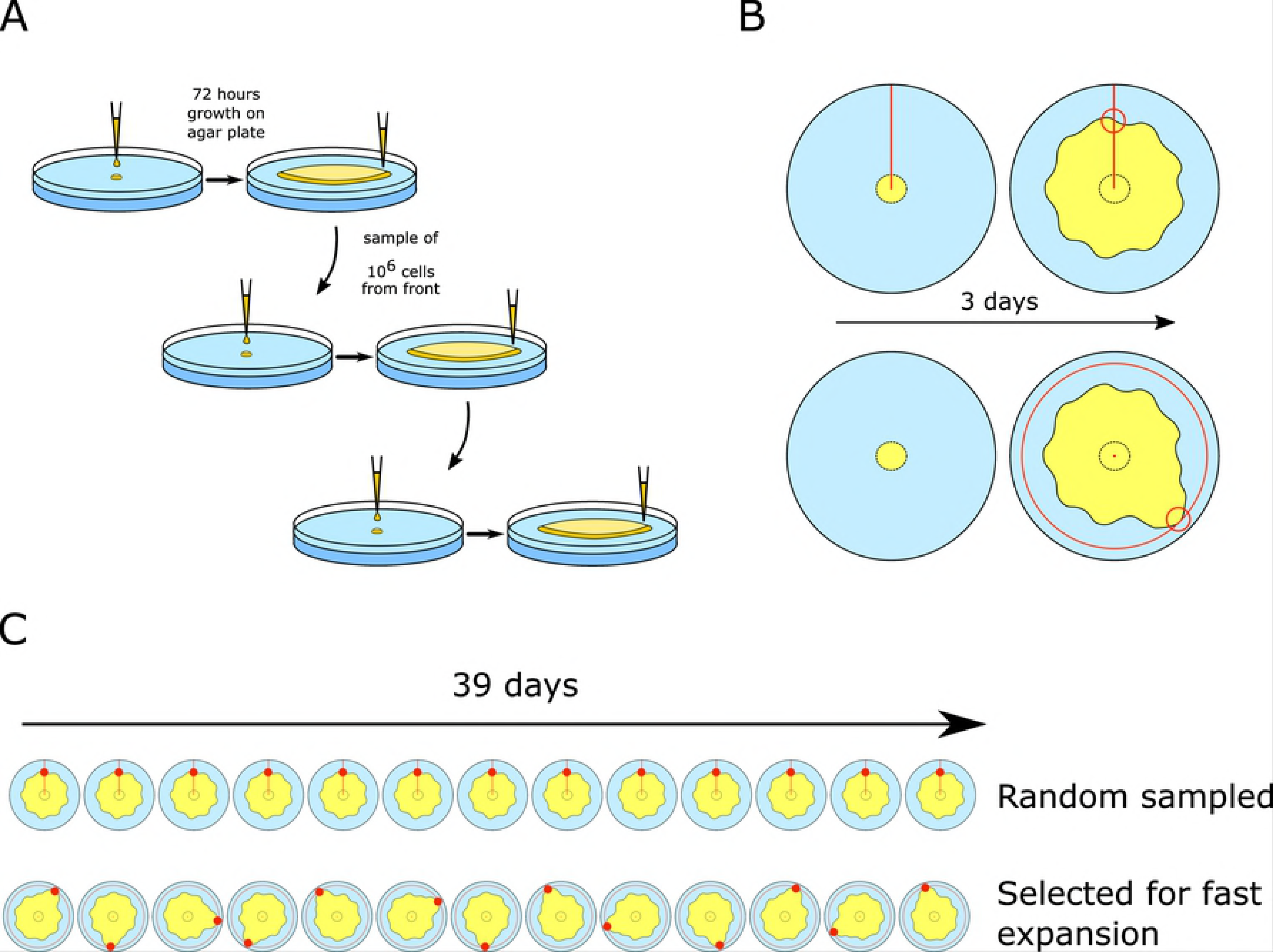
Experimental setup. A) ~100 million bacteria were sampled after 3 days of growth on the edge of the colony and diluted in 100 μl LB medium and one million bacteria were transferred to a new agar plate for a new growth cycle. B) Two sampling protocols were used: No selection CONTROL) we drew a line from the centre of the plate to the edge before the plate was inoculated, and after three days of growth, bacteria were sampled at the point of contact of the line and the colony and artificial selection (SEL): we let the bacteria grow for 3 days, and the edge of the furthest expanding sector of the colony was sampled. C) E. coli lines were grown agar plates for a total of 39 days (1650 generation) with two different sampling protocols: Random sampled and selected for fast expansion.

### DNA extraction

After the range expansion experiment, one million cells from the wave front were streaked out on an LB agar plate containing 0.5% arabinose and incubated for 24h at 37°C to isolate single clones. A single colony was dissolved in 100 μl dilution solution (0.85% NaCl) and 1 μl was transferred to a new LB agar plate containing 0.5% arabinose. The plate was then incubated for 24h at 37°C. Then, the entire colony was removed from the agar plate and resuspended in 1 ml dilution solution. Genomic DNA was extracted using the Wizard Genomic DNA Purification Kit (Promega) following the manufacturer protocol. The integrity of the DNA was checked by gel electrophoresis. The DNA concentration was determined by fluorometric quantification (Qubit 2.0).

### Whole genome sequencing and variant calling

Twenty SEL lines selected for fast expanding speed were sequenced using a TruSeq DNA PCR-Free library (Illumina) on a HiSeq 3000 platform (Illumina), from which we obtained 100bp end reads for all samples. Trimmomatic 0.32 (30) was used to remove the adapter sequences from the reads and for quality trimming. Leading and trailing bases with quality below 3 were removed. The reads were scanned with a 4bp sliding window, and cut if the average quality per base was below 15. Reads with a length below 36 were excluded from the analysis. Variants were identified using BRESEQ, a computational tool for analyzing short-read DNA data (31). The reads were mapped to the *E. coli* K12 MG1655 reference genome (NC_000913.3)

### Estimation of dN/dS ratio

The synonymous and non-synonymous substitutions in each line were counted. The dN/dS ratio was computed by taking the expected number of synonymous and non-synonymous substitutions into account if all codon positions in the reference genome would mutate.

### Expansion velocity on agar plate

Images of the colony were taken during the experiments on agar plates (n=20 for CONTROL lines, n=20 for SEL lines) before transferring the cells to a new plate. We took a picture every three days for each line, and thus have a total of 13 pictures for each line. The images were analyzed with the Fiji package of the imageJ software (32). The radius of the colony was measured and plotted against time. The change in expansion velocity was then determined by fitting a mixed-effect linear model to the data. The observations were arranged in a one-way classification. This means that data were grouped by the 20 samples that were repeatedly measured over time. This model assumes that the growth rate of all lines changes due to a fixed effect *β* common to all lines, but it considers line-specific variability in growth rates by including random effects ***b***_*i*_ for the intercept and slope for the *i*-th line, as:

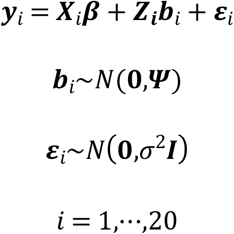

where ***X***_*i*_ and ***Z***_*i*_ are known fixed effect and random effect regressor matrices, ***ε***_*i*_ is the within group error with a Gaussian distribution, and ***ψ*** is the variance-covariance matrix of the random effects. Two nested models were compared with a likelihood-ratio test. The first linear model is defined as

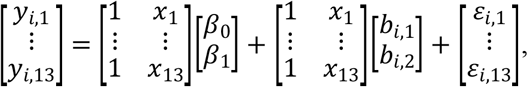

whereas the second model includes an additional quadratic term (*β*_2_) for fixed effects, as

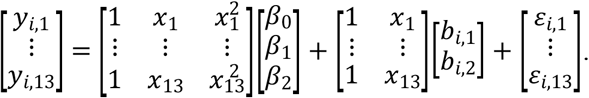

### Determining important genes for expansion speed by shrinkage methods

The mixed effect model from the expansion velocity analysis was used to predict the colony size after 39 days. The data were fist log-transformed as

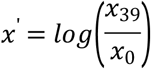

where x_39_ is the predicted colony size at day 39 and x_0_ is the predicted initial colony size at the start of the experiment. A value of 0 indicates that the colony size stays at the same level, positive values indicate an increase, and negative values a decrease in colony size.

For each bacterial line, we counted all non-synonymous, frameshift, and nonsense mutations for each gene. The elastic net method implemented in the R package glmnet was used to fit a Gaussian model to the data. Elastic net is a modification of linear regression that imposes a penalty on the magnitude of the coefficients. Roughly speaking, we performed a linear regression where we forced the estimated parameters to be small via a penalty that is proportional to the sum of the absolute values of the estimated parameters of the linear regression model to avoid overfitting. Avoiding overfitting is particularly important if the number of response variables is large relative to the number of data points. The penalty is controlled by the regularization parameter *λ*, whose value was chosen by 3-fold cross-validation using the cv.glmnet function of the glmnet package. Elastic net is a combination of two model choice methods, namely ridge regression and LASSO (33). The elastic net penalty is controlled by the parameters μ where α = 0 corresponds to LASSO and α = 1 to ridge regression.

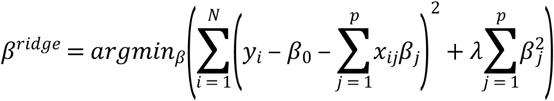

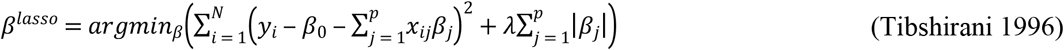

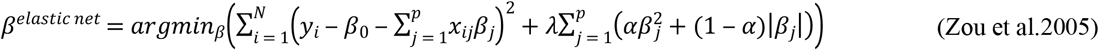

### Distribution of mutations on the bacterial genome

In order to determine signals of convergent adaptation we were searching for mutations that have targeted the same gene in unrelated lines. We calculated the distribution of mutations on the bacterial genome by assignig the 4530 observed SNPs that accumulated in CONTROL lines into 100 equal-sized bins, where each bin was approximately 50 kb in size (**Figure S1**). A Morlet wavelet power spectrum was computed with the *analyze.wavelet* function of the WaveletComp R package. The extracted power levels were used to reconstruct the mutation pattern over the genome by applying the *reconstruct* function of the Wavelet package, and we extracted the probability *P*_*hit*_ (i) that a gene *i* is hit by a mutation. We simulated the same number of mutations as we found non-synonymous substitutions and synonymous substitutions in the CONTROL and SEL lines by generating multinomially distributed random number vectors with a length of the total numbers of genes and the probabilities that we determined by the wavelet analysis. We simulated this vector 1000 times and we calculated the mean value as well as the 2.5% and 97.5% quantiles from the simulated data and compared to the observed CONTROL and SEL data.

### Distribution of simple sequence repeats (SSR) on the bacterial genome

All simple sequence repeats (SSRs) longer than 5 nucleotides were extracted from the *E. coli* K12 MG1655 reference genome (NC_000913.3). For the SSRs that were located in coding regions, the gene of residence was determined. To test whether the SSRs are enriched in specific pathways or functions of the bacterial cell, two subsets of genes were considered: i) a list of genes that do not have any SSR with a length larger or equal 5, and ii) the top 10% of genes with the highest density of SSR. The density of the SSRs in a certain gene was determined by taking the number of SSRs with a length larger or equal to 5 in the gene and dividing it by the length of the gene. A gene ontology term enrichment analysis was performed using the topGO package for R (34).

### Gene ontology enrichment test

In the gene ontology (GO) enrichment analysis, we only used non-synonymous, frameshift and nonsense mutations. The test was performed with the topGO package for R (34). In the GO enrichment analysis, we specifically use information on the number of mutations having occurred in each gene. For any gene the probability from the wavelet analysis that it is hit by a mutation was used to perform a one-tailed binomial test to calculate for any given gene if the observed number of mutations deviated from the expected number. The resulting list of significant genes was used to perform a Fisher’s exact test to determine significantly over-represented GO terms. The *weight01* algorithms used in the topGo analysis iteratively removes the genes mapped to significant GO terms from higher level GO terms and the significance score of connected nodes are compared to detect the locally most significant terms in the GO graph by down-weighting genes in less significant neighbors. Separate analyses were done on CONTROL (n =20) and SEL (n = 20) lines.

### Motility test

20 CONTROL and 20 SEL lines were incubated in LB medium with addition of 0.2 % arabinose at 37°C for 24h. The cultures were diluted to an optical density (OD600) of 0.8 with 0.85% NaCl solution. 2.5 ul of the bacterial suspension was transferred to a LB agar plate with 0.3% agar. The plates were incubated for 18h at 37C°. The tested line is motile if the growth is radiating away from the central inoculation point (35).

## Acknowledgments

We are grateful to Tosso Leeb, Cord Drögemüller and the NGS core facility of the University of Beme for their support.

---

**Figure S1:** Distribution of mutations on the genome in SEL and CONTROL lines. The position of the origin of replication is indicated by a dashed line. Blue: CONTROL lines, Orange: SEL lines, dark red: overlap of CONTROL and SEL lines

**Figure S2:** Genes showing multiple mutations in **A:** SEL lines (red). and **B:** CONTROL lines (blue). The number of mutations in a gene is indicated by the intensity of the color.

**Figure S3:** Motility experiment. Blue: CONTROL lines, Red: SEL lines. The identifier of the CONTROL and the SEL lines are indicated in the plot. The y axis indicates the distance that the lines travelled in 24 h. The dashed gray line indicates the expansion distance of the ancestor line.

**Figure S4:** Coefficients of the elastic net analysis. Blue: genes where a mutation leads to a decrease in colony size. Red: genes where a mutation leads to an increase in colony size.

**Figure S5:** Mosaic Plot of the number of significant genes from the Elastic Net analysis. The gene function group are indicated at the top. Red: Genes where a mutation leads to an increase in colony size. Blue: Genes where a mutation leads to a decrease in colony size. The areas are proportional to the number genes in the group combination.

**Table S1:** Coefficients inferred by elastic net regression. The coefficients represents the effects of mutations on bacterial growth and are relative to the initial colony size. Only non-synonymous mutations were used for the analysis. Orange: genes that have mutations in SEL lines, blue: genes that have mutations in CONTROL lines.

**Table S2:** Coefficients inferred by elastic net regression. The coefficients represents the effects of mutations on bacterial growth and are relative to the initial colony size. Only Frame shift and nonsense mutations were used for the analysis. Orange: genes that have mutations in SEL lines, blue: genes that have mutations in CONTROL lines, violet: genes that have mutations in both CONTROL and SEL lines.

**Table S3:** GO enrichment. Only non-synonymous, frame shift, and nonsense mutations were used for the analysis. The highlighted GO terms represent significant terms after correction for multiple testing (FDR). Orange: SEL lines. Blue: CONTROL lines.

**Table S4:** Mutations in SEL and CONTROL lines. An asterisk (*) indicates genes associated with the “taxis” GO term. Highlighted genes are mutated in both conditions.

**Table S5:** GO enrichment analysis using the top 10% of genes with the highest density of SSRs with a length larger or equal 5.

**Table S6:** GO enrichment analysis with genes that do not have any SSRs with a length larger or equal

